# LDL exposure disrupts mitochondrial function and dynamics in a hippocampal neuronal cell line

**DOI:** 10.1101/2024.06.12.598647

**Authors:** Hémelin Resende Farias, Jessica Marques Obelar Ramos, Caroline Griesang, Lucas Santos, Osmar Vieira Ramires Junior, Debora Guerini de Souza, Fernanda Silva Ferreira, Sabrina Somacal, Leo Anderson Meira Martins, Diogo Onofre Gomes de Souza, José Cláudio Fonseca Moreira, Ângela Terezinha de Souza Wyse, Fátima Theresinha Costa Rodrigues Guma, Jade de Oliveira

**Author notes:** Correspondence: Jade de Oliveira, Programa de Pós-Graduação em Ciências Biológicas: Bioquímica, Departamento de Bioquímica, Instituto de Ciências Básicas da Saúde, Universidade Federal do Rio Grande do Sul (UFRGS), Porto Alegre, Brazil.

## Abstract

Hypercholesterolemia has been associated with cognitive dysfunction and neurodegenerative disease. Moreover, this metabolic condition disrupts the blood-brain barrier, allowing Low-Density Lipoprotein (LDL) to enter the Central Nervous System. Thus, we investigated the effects of LDL exposure on mitochondrial function in a mouse hippocampal neuronal cell line (HT-22). HT-22 cells were exposed to human LDL (50 and 300 μg/mL) for 24 hours. After this, intracellular lipid droplet (LD) content, cell viability, cell death, and mitochondrial parameters were performed. We found that the higher LDL concentration LDL increases LD content compared to control. Both concentrations increased the number of Annexin V-positive cells, indicating apoptosis. Moreover, in mitochondrial parameters, the exposure of LDL on hippocampal neuronal cell line leads to a decrease in mitochondrial complexes I and II in both concentrations tested and a reduction in Mitotracker™ Red fluorescence and Mitotracker™ Red and Mitotracker™ Green ratio in the higher concentration, indicating dysfunction in the mitochondria. The LDL incubation induces mitochondrial superoxide production and a decrease in superoxide dismutase activity in the lower concentration in HT-22 cells. Finally, hippocampal neuronal cell line exposed to LDL exhibit an increase in the expression of genes associated with mitochondrial fusion (OPA1 and Mitofusin 2) in the lower concentration. In conclusion, our findings suggest that LDL exposure induces mitochondrial dysfunction and modulation in mitochondrial dynamics in the hippocampal neuronal cells.

## 1 INTRODUCTION

Hypercholesterolemia is a metabolic disorder characterized by high levels of plasmatic cholesterol [1]. It is already established that elevated plasmatic cholesterol levels are risk factors for developing atherosclerotic cardiovascular disease and stroke [2]. Moreover, in the last decades, hypercholesterolemia has been associated with the development of cognitive impairments characteristic of neurodegenerative diseases, such as Alzheimer’s disease [3–6]. The lipoprotein more associated with health prejudice caused by hypercholesterolemia is the low-density lipoprotein (LDL)[7]. This lipoprotein is responsible for transporting cholesterol from the blood to the tissues [8, 9] ; however, it is easily oxidized in peripheral tissues[10].

The exact mechanism by which hypercholesterolemia leads to neuronal damage and, consequently, cognitive impairment is unclear, but some mechanisms have been proposed. Experimental studies in hypercholesterolemic rodents demonstrated alterations in the cholinergic system [11–13], reduced mitochondrial metabolism [13–16], increases in reactive species production [13–17] and alterations in antioxidant enzymes activity [14, 15, 17, 18] in different brain regions.

Cholesterol metabolism in Central Nervous System (CNS) occurs independently of peripheral metabolism since the plasma lipoproteins cannot cross the blood-brain barrier (BBB) [19]. Therefore, under normal conditions, brain cholesterol is derived from astrocytes [20, 21]. However, hypercholesterolemia is associated with BBB disruption [13, 22, 23], as well as neuroinflammation [11, 13, 24, 25], therefore allowing the entry of compounds of the peripheral system to the CNS, such as LDL and inflammatory factors [26, 27].

*In vitro* studies have already demonstrated that LDL causes an increase in reactive species production in human neuroblastoma cells [28], disturbs the structure and function of endolysosomes, as well as increases the Aβ production in primary neurons [29]. However, the direct effect of LDL on mitochondria from neurons is unclear. The neurons have high energetic demands to exercise their functions and require that their mitochondria are viable [30–33]. The mitochondria are responsible for several metabolic functions, such as ATP production. Moreover, the mitochondria are the primary source of reactive species production, mainly in oxidative phosphorylation [34]. Thus, when something causes mitochondrial dysfunction, especially mitochondrial complexes alteration, it leads to energetic deficits and oxidative stress [35].

Mitochondria are remarkably dynamic organelles that control their size, morphology, and number through fusion and fission, these processes are known as mitochondrial dynamics [36]. The maintenance of the mitochondrial organization, function, and morphology is complex, and it is orchestrated by a group of proteins that maintain the equilibrium between form and function by coordinating their activities [37]. These proteins are dynamin-related protein1 (Drp1), mitofusin (Mnf) 1 e 2, and Optic Atrophy 1 (OPA1). For mitochondrial fusion, the main proteins associated are Mnf 1 e 2, which mediate outer mitochondrial membrane fusion, while OPA1 mediates the inner mitochondrial membrane fusion [36]. The main protein associated with mitochondrial fission is Drp1 [38]. Therefore, herein, we investigated the effects of LDL exposure in HT-22 cells, mainly on mitochondrial function.

## 2 MATERIALS AND METHODS

### 2.1 Cell Culture and Experimental Design

HT-22 cells (Mouse Hippocampal Neuronal cell line) were grown in Dulbecco’s modified Eagle’s medium (DMEM, Sigma, D7777) containing 10% fetal bovine serum (CRIPION, SP, Brazil) and 100 IU penicillin/streptomycin (Sigma, P0781), at 37 °C in 5% CO_2_ and 95% air in a humidified atmosphere. HT-22 cells were seeded at the density of 2x10^4^/cm^2^, and after 24 hours of growing, HT-22 cells were exposed to LDL (50 or 300 μg/ml) for 24 hours. The concentrations were chosen from previous experiments that observed alterations caused by LDL exposure in neurons [28, 29]. All experiments were repeated at least three times. LDL was isolated from normolipidemic human serum by discontinuous density-gradient ultracentrifugation in KBr solutions containing 30 mmol/l EDTA as described by de Bem et al [39]. The concentration of LDL was determined from the total protein concentration, and protein content was quantified by the method described by Lowry et al. [40], using bovine serum albumin as standard. Ethical Committee project approved under number 4557728.

### 2.2 AdipoRed™ assay

Intracellular lipid droplets were quantified using the AdipoRed™ Assay Reagent (Lonza, PT-7009) according to the manufacturer’s protocol. Briefly, after LDL exposure, the cells were prewashed with PBS once and incubated with the AdipoRed™ Reagent for 15 min (1:40). Data acquisition and analysis were performed in FACSCalibur™ flow cytometer (BD Bioscience, San Jose, CA, USA). Data were analyzed using FlowJo XV version 10 (FlowJo LLC)

### 2.3 Cell viability assay

The cell viability was assessed by the reduction of 3-(4,5-Dimethylthiazolyl)-2,5-diphenyltetrazolium bromide (MTT) assay. After LDL exposure, we added a MTT (Sigma, M2128) solution (5 mg/mL dissolved in PBS sterile) to the medium, reaching a final concentration of 0.5 mg/mL. The cells were incubated for three hours at 37ºC, and the formazan formed by the reduction of MTT was dissolved in dimethyl sulfoxide (DMSO). Results are expressed as the percentage of control cells. All experiments were performed in technical triplicate. This protocol was adapted from Farias et al [41].

### 2.4 Cell Density

To assess cell density and determine the effects of LDL exposure on cell survival, we utilized the sulforhodamine B (SRB) assay. After the LDL exposure, HT-22 cells were stained with 0.4% sulforhodamine B (Sigma, S1402) in acetic acid 1% for one hour at room temperature. Excess unbound SRB was removed by washing the cells five times with distilled water. The stained cells were then dissolved in 1% SDS, and the absorbance was measured at 560 nm using the SpectraMax® M5 (Molecular Devices). Results were expressed as a percentage of control.

### 2.5 Lactate dehydrogenase (LDH) activity

LDH release was performed to test the loss of plasma membrane integrity. After LDL exposure, the culture medium was collected, centrifuged, and analyzed. The LDH activity was performed using the LDH diagnostic kit according to the manufacturer’s instructions (BioTecnica).

### 2.6 AnnexinV positive-cells

The FITC Annexin V (QuatroG, 100034) was performed for cell death analysis following the manufacturer’s instructions. Samples were incubated in a binding buffer containing Annexin-V FITC and PI for 15 minutes in the dark at room temperature. As a positive control, cells were frozen and then stained. Data acquisition and analysis were performed using a FACSCalibur™ flow cytometer (BD Bioscience, San Jose, CA, USA). Data were analyzed using Data were analyzed using FlowJo XV version 10 (FlowJo LLC).

### 2.7 Mitochondrial Complexes activities

Mitochondrial complex I (NADH dehydrogenase) activity was measured by the NADH-dependent ferric reduction rate at 420 nm, as described by Cassina and Radi [42]. The activity was calculated in nanomoles per minute per milligram of protein. The complex II activity was measured according to Fischer et al [43] by the decrease in absorbance due to the reduction of 2,6-dichloroindophenol (DCIP). The results were expressed as nanomoles per minute per milligram of protein.

### 2.8 Mitotracker™ Red and Green

Mitochondrial mass and membrane potential were evaluated by Mitotracker™ Green (MTG - Invitrogen™, M7514) and Red (MTR-Invitrogen™, M7512) dye, respectively. Therefore, it was possible to establish a relationship between MTR and MTG fluorescence to estimate the rate of mitochondrial function [44]. After LDL exposure, HT-22 cells were harvested using trypsin. Then, cells were resuspended and incubated for 20 minutes in the dark with 100 nM of MTG and 100 nM of MTR diluted into pre-warmed (37 °C) Hank’s Balanced Salt Solution (HBSS). The samples were analyzed using a FACSCalibur™ flow cytometer (BD Bioscience, San Jose, CA, USA) and data were analyzed using FlowJo XV version 10 (FlowJo LLC).

### 2.9 MitoSOX™

For quantitation of mitochondrial superoxide generation, cells were loaded with MitoSOX™ Red (Invitrogen™, #M36008). After LDL exposure, the cells were prewashed with PBS once loaded with Mitosox Red (500nM) in HBSS for 30 min. Cells were then washed. Fluorescence intensity was then measured at 510/580 nm in SpectraMax® M5 (Molecular Devices)[45].

### 2.10 Superoxide dismutase (SOD) activity

SOD activity was determined using the RANSOD kit (Randox, SD125). This method employs xanthine and xanthine oxidase to generate superoxide radicals that react with 2-(4-iodophenyl)-3-(4-nitrophenol)-5-phenyltetrazolium chloride (INT) to form a formazan dye that is assayed by spectrophotometric analysis at 505 nm at 37 °C in lysed cells. SOD activity is expressed as U/mg of protein. One unit of SOD causes a 50 % inhibition of the rate of reduction of 2-(4-iodophenyl)-3-(4-nitrophenol)-5-phenyltetrazolium chloride under the conditions of the assay.

### 2.11 Gene expression analysis (RT-qPCR)

RNA extraction was performed using TRIzol® Reagent (ThermoFisher Scientific, USA) and 2-Mercaptoethanol (Sigma-Aldrich, M3148) following the manufacturer’s protocol and Santos et al. [46]. RNA concentration and purity were quantified using the I-Quant equipment (Loccus, BR), with purity verified through the ratio of absorbances at 260nm and 280nm (A260/A280). The cDNA synthesis reaction was then performed using 2 μg of RNA for each sample with the High-Capacity cDNA Reverse Transcription® kit (Thermo Fisher Scientific, USA).

Gene-specific primers were designed using IDT Design software (Integrated DNA Technologies Inc., USA), ensuring no secondary structures were generated. Primer efficiency was evaluated to confirm the absence of nonspecific amplifications. Subsequently, gene expression analysis for proteins involved in mitochondrial dynamics: Mitofusin 1 (MFN1), mitofusin 2 (MFN2), optic atrophy protein 1 (OPA1), and dynamin-related protein 1 (DRP-1) and β-actin housekeeping gene was conducted using the sequences shown in Table 1. RT-qPCR reactions were performed in triplicate using the PowerUp™ SYBR® Green Master Mix kit (Thermo Fisher Scientific, USA), following the manufacturer’s instructions. The results were analyzed using the 2−ΔΔCT method.

**Table 1.**
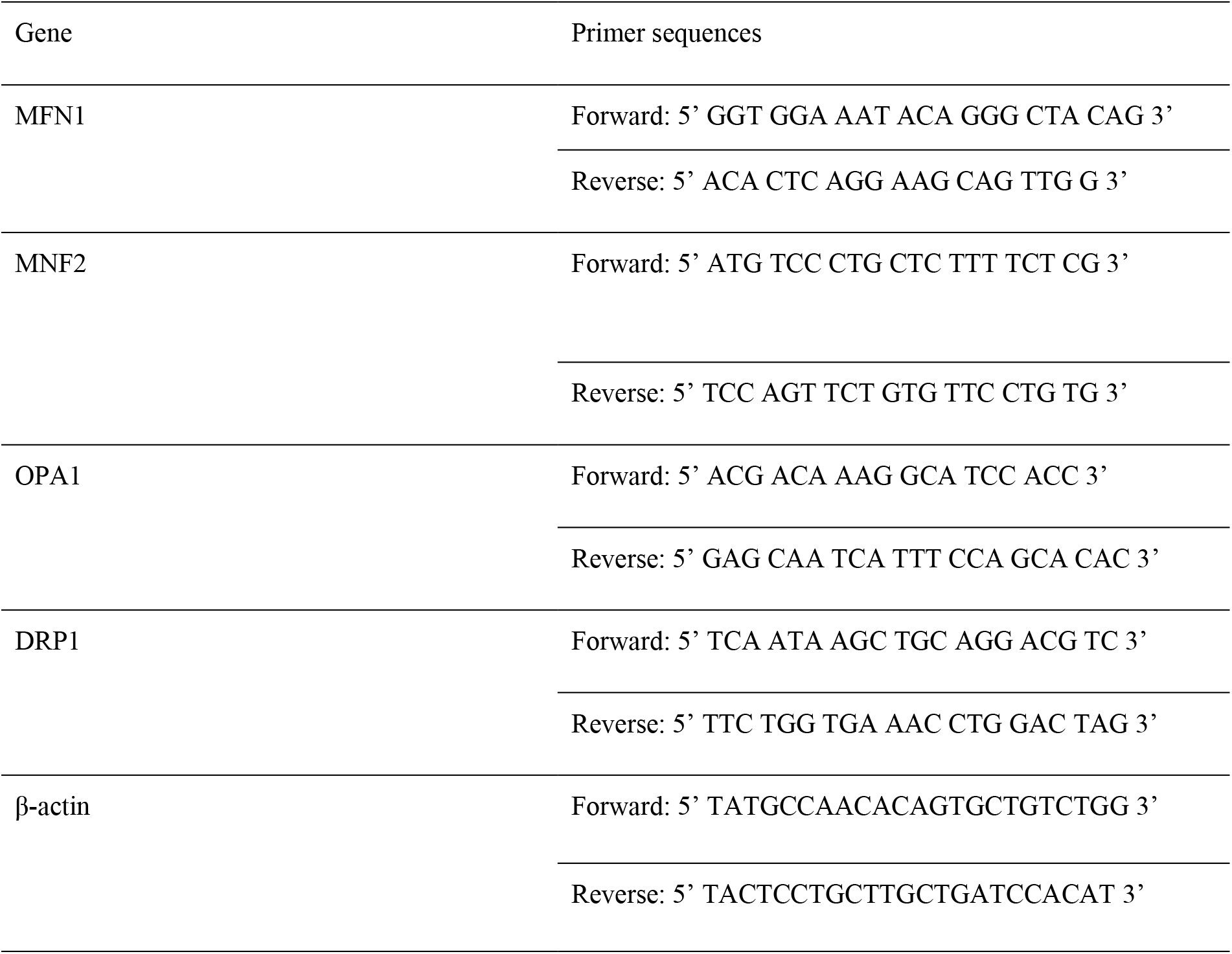
Sequence of primers used for RT-qPCR.

### 2.12 Statistical analyses

All experiments were performed on at least three occasions. Results were expressed as mean ± standard error of the mean (SEM). One-way ANOVA with Dunnet post hoc tests was used for data analysis, and a significant difference was defined as p < 0.05.

## 3 RESULTS

### 3.1 LDL exposure leads to an increase in intracellular lipid droplet content in the hippocampal neuron cell line

First, we evaluated the intracellular lipid droplet content after 24 hours of LDL exposure. We observed an increase in the intracellular lipid droplet content at the higher concentration of LDL (Fig. 1a and 1b), confirming the LDL uptake by HT-22 cells.

**Fig. 1.**
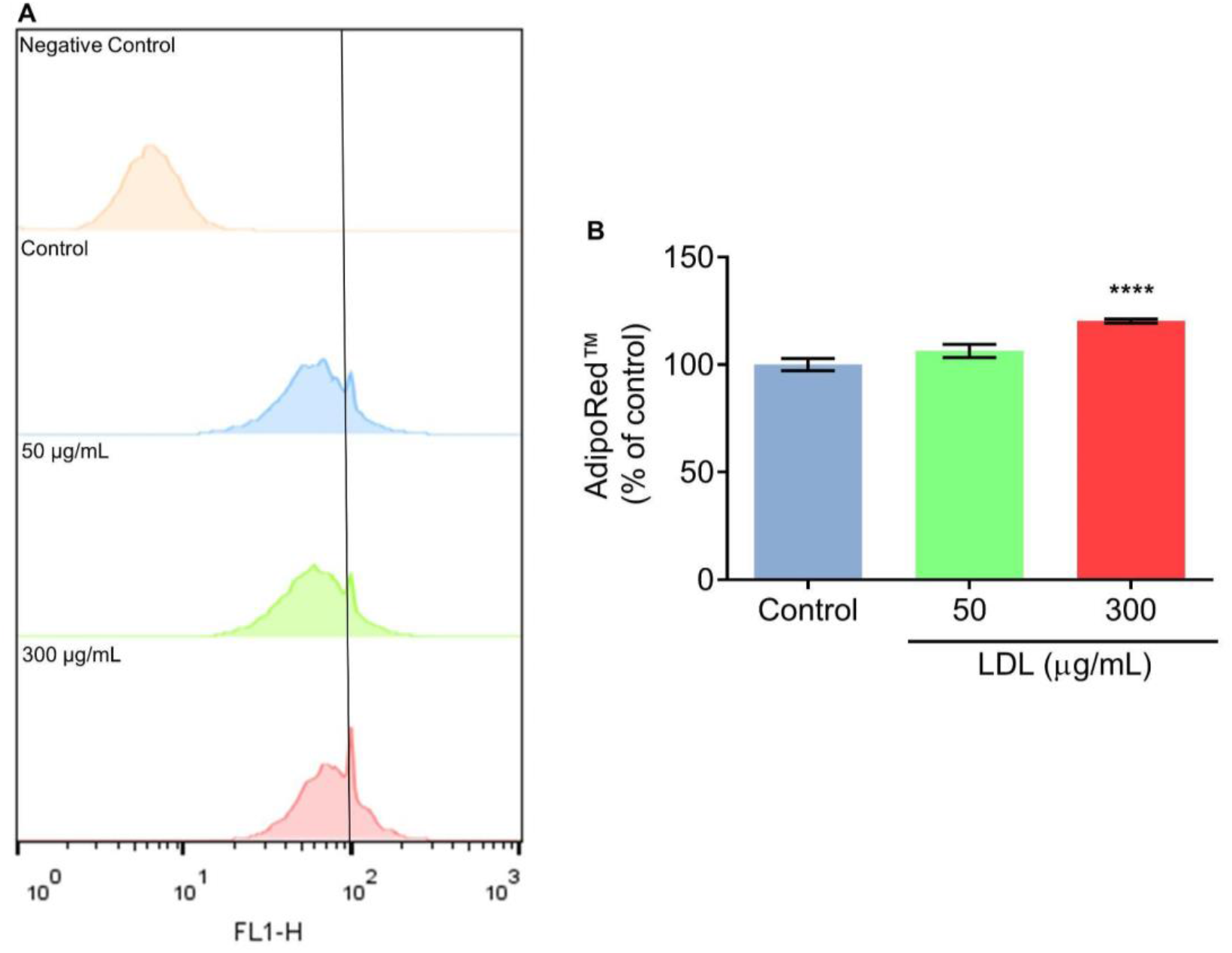
LDL exposure increases lipid droplets in HT-22 cells. (a) Representative histogram of lipid droplets assessed by AdipoRed™. (b)Lipid Droplets. Data were expressed as mean ± SEM. Statistical analyses were performed using a One-way ANOVA/Dunnett post hoc test. ****p<0.0001 compared to control group.

### 3.2 LDL exposure induces an increase in AnnexinV-positive cells in the hippocampal neuron cell line

Exposure to LDL did not affect the viability of hippocampal neuronal cell line, as assessed by the MTT (Fig. 2a) and SRB assays (Fig. 2b). However, when we measured the cellular death by Annexin, we observed that LDL exposure (50 and 300 ug/ mL) leads to a significant increase of AnnexinV-positive cells in HT-22 cells (Fig. 2 d and e), suggesting that LDL induces apoptosis. Moreover, HT-22 cells exposed to LDL have no alteration in extracellular LDH activity, a parameter for necrosis (Fig. 2c).

**Fig. 2.**
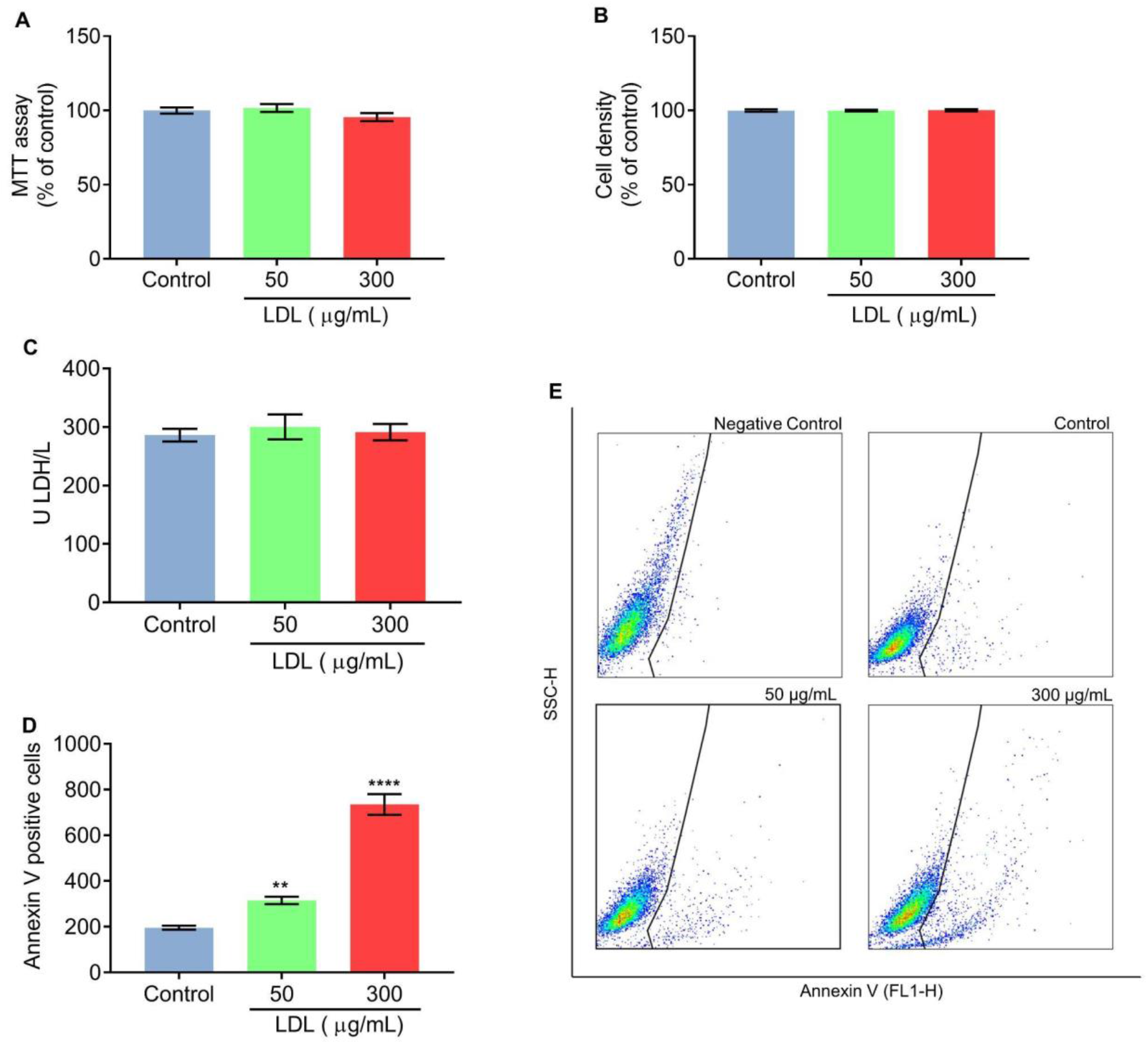
LDL induces an increase in AnnexinV-positive cells in HT-22 cells. (a)The Cell viability was measured using an MTT assay. (b) SRB assay was used to measure cell density. (c) Extracellular LDH activity. (d) and (e) AnnexinV-positive cells, (d)graphic analysis, (e) Dot plot. Data were expressed as mean ± SEM. Statistical analyses were performed using a One-way ANOVA/Dunnett post hoc test. **p<0.01, ****p<0.0001 compared to control group.

### 3.3 LDL exposure leads to a reduction in mitochondrial complexes activities and alteration in mitochondrial membrane potential in HT-22 cells

The effect of LDL on the mitochondria of the hippocampal neuron cell line was evaluated. We observed that the exposure to LDL leads to a decrease in mitochondrial complexes I (Fig. 3a) and II (Fig. 3b) in both concentrations (50 and 300 µg/mL) in HT-22 cells. Afterward, the measurement of mitochondrial mass (MTG) and potential (MTR) was made by flow cytometry. We observed that LDL induces a significant reduction in MTR fluorescence in the concentration of 300 µg/mL (Fig. 3d) without altering the mitochondrial mass (Fig. 4e). We also observed that the exposure to LDL at 300 µg/mL decreases significantly the MTR/MTG ratio (Fig. 3f). Figure 3C shows a fluorescence shift to the left and down upon exposure to 300 µg/mL of LDL in HT-22 cells, indicating mitochondrial damage induced by LDL.

**Fig. 3.**
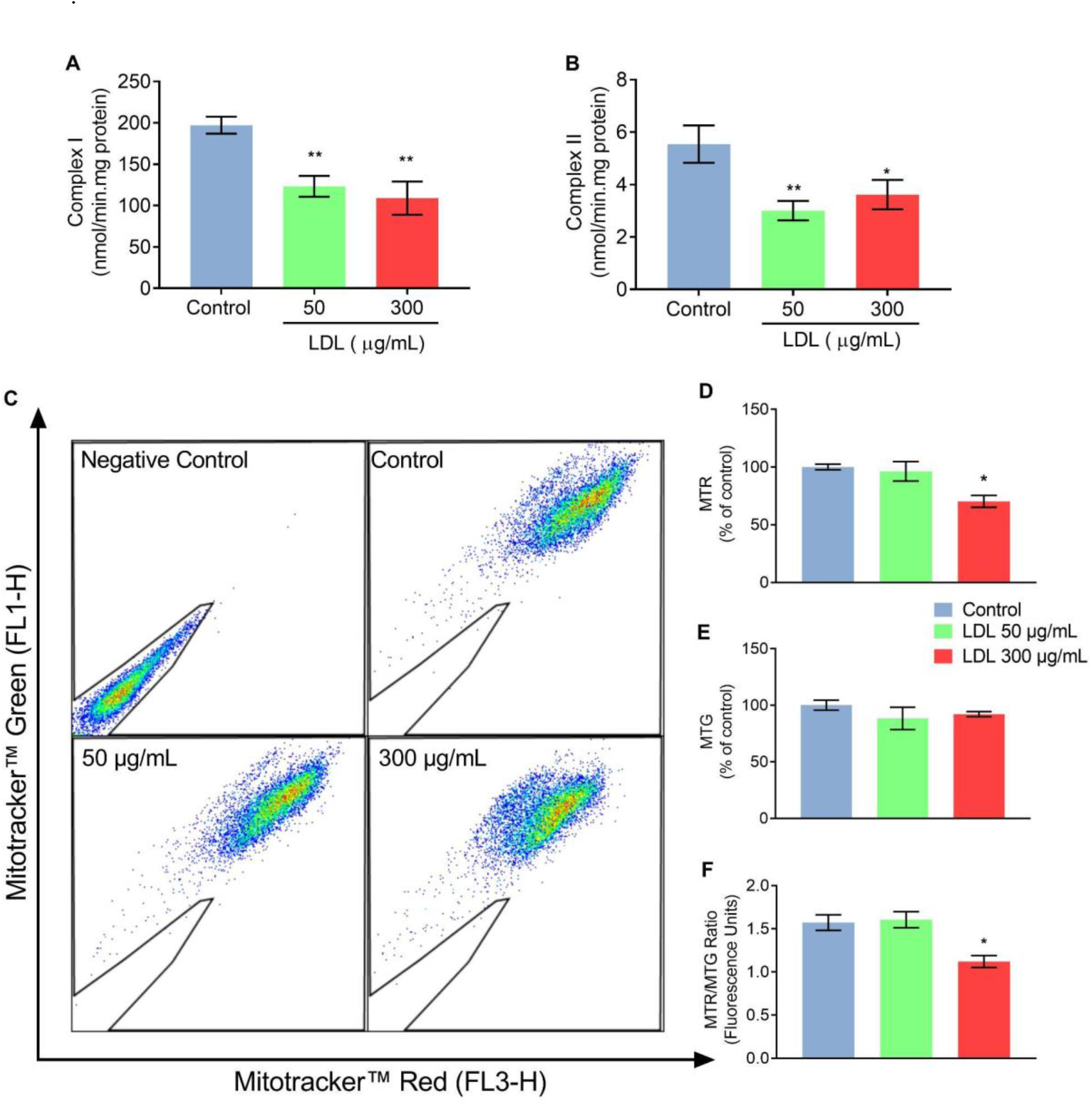
LDL induces mitochondrial dysfunction in the hippocampal neuron cell line. (a) Mitochondrial complex I activity. (b) Mitochondrial complex II activity. (c) Dot plot of Mitotracker™ Red and Green. (d) Graphic representation of Mitotracker™ red fluorescence (% of control). (e) Graphic representation of Mitotracker™ green fluorescence (% of control). (f) Mitotracker™ Red and Mitotracker™ green ratio. Data were expressed as mean ± SEM. Statistical analyses were performed using a one-way ANOVA/Dunnett post hoc test. **p<0.01, ****p<0.0001 compared to control group. Legend: MTR - Mitotracker™ Red; MTG - Mitotracker™ Green

**Fig. 4.**
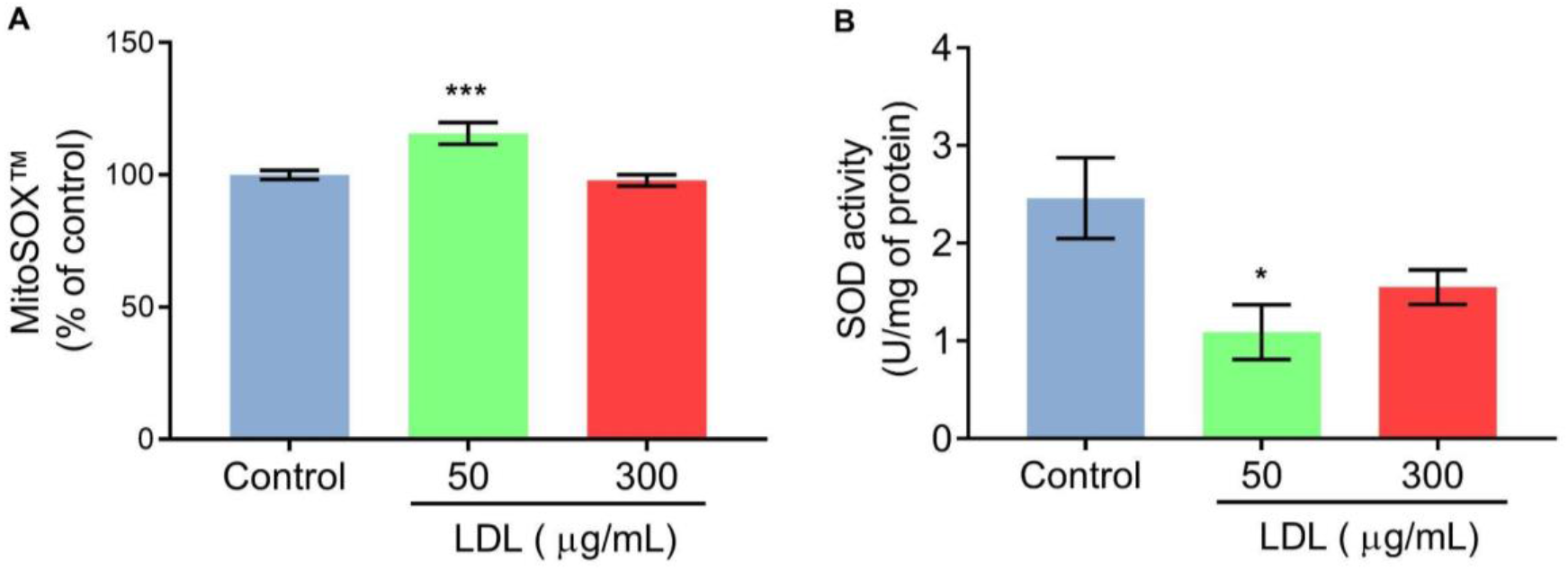
LDL exposure induces an oxidative environment in the hippocampal neuron cell line. (a)Mitochondrial superoxide production assessed by MitoSOX™. (b) Superoxide Dismutase (SOD) activity. Data were expressed as mean ± SEM. Statistical analyses were performed using a one-way ANOVA/Dunnett post hoc test. *p<0.05, ***p<0.001 compared to control group.

### 3.4 LDL exposure induces an increase in the production of mitochondrial superoxide production and reduces superoxide dismutase activity in HT-22 cells

It has been established that mitochondrial dysfunction increases reactive species production [47]. Then, we evaluated the effect of LDL on mitochondrial superoxide production and Superoxide Dismutase activity. Our results demonstrated that the lower concentration (50 ug/mL) of LDL induces a significant increase in mitochondrial superoxide production (Fig. 4a), as well as a significant decrease in Superoxide dismutase activity (Fig. 4b). These results indicated that LDL induces an oxidative environment in HT 22 cells.

### 3.5 LDL exposure induces an increase in the expression of genes associated with mitochondrial fusion in HT-22 cells

We evaluated the effect of exposure to LDL on the expression of genes that code for proteins associated with mitochondrial dynamics. We observed that at the lower concentration of LDL (50 ug/mL), there was a significant increase in the gene expression of OPA1 (Fig. 5B), a protein associated with inner mitochondrial membrane fusion, and MFN2 (Fig. 5d), a protein associated with outer mitochondrial membrane fusion [48]. The LDL exposure did not significantly change the DRP1 and MNF1 expression.

**Fig. 5.**
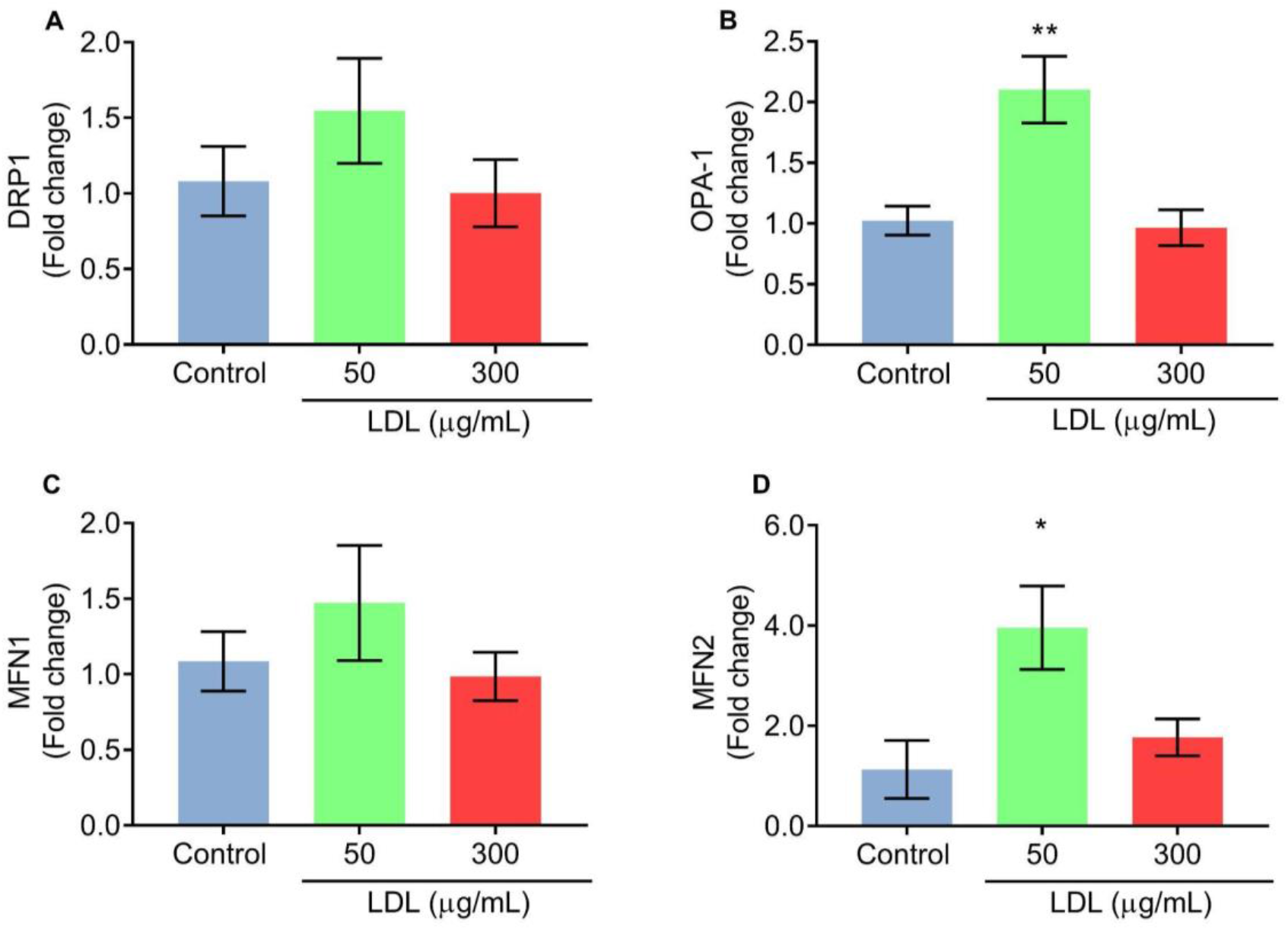
LDL exposure increases the expression of genes associated with mitochondrial fusion in the hippocampal neuron cell line. (a) DRP1 expression, (b) OPA-1 expression. (c) MFN1 expression. (d) MFN2 expression. Data were expressed as mean ± SEM. Statistical analyses were performed using a one-way ANOVA/Dunnett post hoc test. *p<0.05, **p<0.01 compared to control group.

## 4 DISCUSSION

Hypercholesterolemia has been experimentally linked with BBB breakdown and cognitive impairment. [13, 22, 23]. Importantly, the BBB dysfunction dysfunction allows the leakage to brain parenchyma of peripheral system compounds, including LDL-cholesterol [26]Moreover, the hippocampus, a brain area critical for memory, is particularly vulnerable to damage induced by hypercholesterolemia [13, 17, 24, 49]. However, the exact mechanism by which high LDL cholesterol levels lead to neuronal damage and subsequent dementia remains unclear. We hypothesize that neurons exposed to LDL will have impaired mitochondrial homeostasis, which may lead to the activation of cell death mechanisms.

Using a Hydrophilic Stain Nile Red (AdipoRed™), we verified that in the higher concentration of LDL, the intracellular lipid droplet was increased in HT-22 cells. The Nile red stains neutral and polar lipids, including triacylglycerol and cholesteryl esters[50, 51]. In health conditions, neurons do not present high lipid droplet contents since they have a lower capacity to use lipids for energy production[52]. However, lipid droplets are increased in neurons in several neurodegenerative diseases [53–56], and it is associated with cognitive impairment [57].

Next, the cellular viability assay showed that LDL does not alter the cell viability of HT-22 cells assessed by the MTT assay. Similarly, it was previously demonstrated that LDL does not alter this parameter in HT-22 cells (100 μg/ml) using the CCK-8 assay [58], nor in hippocampal neural precursor cells (25 to 200μg/ml), performed by the resazurin assay [59]. Similar experiments using the MTT assay on SHSY-5Y cells (neuroblastoma cells) and embryonic cortical neurons were performed by Dias et al [60] and Sugawa et al. [61], respectively. Dias et al [60] demonstrated that LDL does not alter the MTT assay in different concentrations (0.8, 1.6, 4.0, 8.0 μg/ml) and times of exposure (2 and 16h). The concentrations tested by Dias et al. [60] were significantly lower than those used in this work. However, Sugawa et al. [61] demonstrated that, after 24 hours, 100μg/ml of LDL did not alter MTT assay results in embryonic cortical neurons.

We also observed that LDL causes no alteration in the necrosis parameter in HT-22 cells, assessed by extracellular LDH activity. While LDL does not alter cellular viability or necrosis levels, we observed an increase in Annexin V staining, suggesting apoptosis of HT-22 cells in both concentrations. Annexin V is a dye-labeled phosphatidylserine (PS)-binding protein [62]. PS is localized exclusively on the inner leaflet of the cell membrane [63, 64]. After induction of apoptosis, PS is flipped to the outer membrane and acts as an “eat me” signal to recruited phagocytes [65–67]. This increase in Annexin V fluorescence can indicate that LDL is leading to apoptosis. However, it is important to mention that Annexin V staining alone is not sufficient to definitively confirm cell death since the plasmatic membrane can be intact. A previous study using primary cortical neurons of rats observed TUNEL-positive staining after exposure to 100 μg/ml LDL, suggesting apoptosis [61]. In contrast, Engel et al [59] did not find evidence of pyknotic nucleus presence, another indicator of apoptosis, in hippocampal neural precursor cells treated with different concentrations of LDL (25 to 200 μg/ml). These findings suggest that LDL can induce neurotoxicity and possible apoptosis in some neuron cells, but further studies are needed to confirm this hypothesis.

Neurons require properly functioning mitochondria to maintain their activity, as they have high ATP demands [68]. Mitochondria have several metabolic functions, mainly combining electron transport along the electron transport chain with oxygen consumption and generation of ATP [69]. These organelles are also strictly involved with apoptosis [70, 71]. Our previous work using hypercholesterolemic animals pointed out a negative correlation between cholesterol levels and complex I and II activities in the cerebral cortex [14]. Moreover, Paul and Borah [13] demonstrated a reduction in complex I and II activities in different brain areas, including the hippocampus, in an experimental model of hypercholesterolemia. Consistent with these findings, our current data demonstrate that LDL exposure in HT-22 cells caused a decrease in both complex I and II activities, suggesting that LDL impairs the activity of these respiratory chain complexes.

To better understand the effect of LDL on mitochondria, we assessed the mass and mitochondrial potential with Mitotracker™ Green and Red, respectively, using flow cytometry. MTR gets across the cellular membrane and accumulates in active mitochondria, depending on their oxidative activity. In addition, MTG labels mitochondria independently of the membrane potential, providing a readout related solely to mitochondrial mass. Therefore, by performing the ratio between MTR and MTG, it is possible to establish the rate of mitochondrial function [72, 73]. Our data suggests that exposure to a higher concentration of LDL (300ug/mL) in HT-22 cells was associated with a reduction in MTR fluorescence. Nonetheless, LDL at 300 μg/ml reduced the MTR/MTG ratio in the neurons, suggesting a possible mitochondrial alteration. The reduction in the MTR/MTG ratio can suggest a mitochondrial swelling, which, together with a decrease in mitochondrial membrane potential, may be correlated with neuronal death [74–76].

It has been known that impairment of mitochondria caused by decreases in the mitochondrial complex is responsible for producing reactive species, such as superoxide anions [77–80]. Here, we found that LDL (50μg/ml) exposure increases mitochondrial superoxide anion production in HT-22 cells. Moreover, a previous study demonstrated in other brain cells, such as astrocytes and microglia cultures, that LDL causes a significant increase in reactive species generation [81], as well in human neuroblastoma cells (SHSY-5Y) [28]. It is important to mention that we observed that LDL induces a decrease in complex I and II activity in both concentrations tested, however, there was an increase in mitochondrial superoxide production only in the cells exposed to 50 μg/mL of LDL. It can be explained by some antioxidant modulation, mainly for Superoxide Dismutase (SOD) activity. We observed that at 50ug/mL, LDL leads to a decrease in SOD activity, suggesting that superoxide production is increased at this exposure concentration. It was in agreement with previous studies that have demonstrated that hypercholesterolemia leads to a decrease in SOD activity in the hippocampus of rodents [82–84].

Hypercholesterolemia has been associated with oxidative stress in both peripheral and central nervous systems [14, 15, 17, 85]. An oxidative environment, characterized by high levels of ROS, is associated with changes in mitochondrial dynamics [86], including an increase in mitochondrial fusion [87]. Our results demonstrated that the lower concentration of LDL leads to an increase in the expression of OPA-1 and Mnf2 genes, both associated with mitochondrial fusion [48]. Mitochondrial fusion occurs when mitochondria merge their outer and inner mitochondrial membranes, resulting in mitochondrial elongation. Fusion facilitates the distribution and mixing of mtDNA, metabolites, proteins, and lipids, acting as a protective mechanism against partially dysfunctional mitochondria by diluting damaged components [88].

In summary, the lower concentration of LDL triggered an increase in mitochondrial superoxide production and a decrease in antioxidant defense. This increase in reactive species appears to induce a modulation in mitochondrial fusion as evidenced by the increased expression of genes associated with fusion at this LDL concentration. Interestingly, the lower LDL concentration did not alter the MTR/MTG ratio. This might be explained by the modulation of mitochondria fusion, potentially preserving the mitochondrial membrane potential [89]. Our findings suggest a link between hypercholesterolemia and cognitive impairment, possibly mediated by LDL-induced mitochondrial dysfunction. Notably, we demonstrate that not only oxidized LDL but also the non-oxidized form of lipoprotein, can cause neuronal damage.

Nevertheless, further investigation is essential to definitively determine the impact of LDL on neuronal death pathways, as well as mitochondrial images, to confirm mitochondrial fusion. Furthermore, the potential influence of LDL on microglia and astrocytes, and the subsequent effects of mediators released by these glial cells on neurons, cannot be disregarded and warrant further exploration.

## Fundings

This study was supported by the Federal University of Rio Grande do Sul (UFGRS), Conselho Nacional de Desenvolvimento Científico e Tecnológico (CNPq) [CNPq/MCTI/FNDCT No 18/2021 - Faixa A - Grupos Emergentes No do Processo: 407006/2021-4], Coordenação de Aperfeiçoamento de Pessoal de Nível Superior (Capes), Fundação de Amparo à pesquisa do Estado do RS [21/2551-0000 740-0 Edital FAPERGS | 10/2020 Auxílio Recem Doutor -ARD], Brazilian National Institute of Science and Technology on Excitoxicity and neuroprotection (INEN 2014-465671/2014-4) and Brazilian National Institute of Brain Health (INSC 406020/2022-1).

## Notes

### Competing Interest Statement

The authors have declared no competing interest.

## References

1. Martinez-Hervas S, Ascaso JF (2023) Hypercholesterolemia. Encycl Endocr Dis 320–326. 10.1016/B978-0-12-801238-3.65340-0

2. Mozaffarian D, Benjamin EJ, Go AS, et al (2015) Heart disease and stroke statistics--2015 update: a report from the American Heart Association. Circulation 131:e29–e39. 10.1161/CIR.0000000000000152

3. Kivipelto M, Helkala EL, Laakso MP, et al (2001) Midlife vascular risk factors and Alzheimer’s disease in later life: Longitudinal, population based study. Br Med J. 10.1136/bmj.322.7300.1447

4. Kivipelto M, Solomon A (2006) Cholesterol as a risk factor for Alzheimer’s disease - epidemiological evidence. Acta Neurol Scand 114:50–57. 10.1111/j.1600-0404.2006.00685.x

5. Zambón D, Quintana M, Mata P, et al (2010) Higher incidence of mild cognitive impairment in familial hypercholesterolemia. Am J Med. 10.1016/j.amjmed.2009.08.015

6. de Bem AF, Krolow R, Farias HR, et al (2021) Animal Models of Metabolic Disorders in the Study of Neurodegenerative Diseases: An Overview. Front Neurosci 14:604150. 10.3389/FNINS.2020.604150/BIBTEX

7. Benito-Vicente A, Uribe KB, Jebari S, et al (2018) Familial Hypercholesterolemia: The Most Frequent Cholesterol Metabolism Disorder Caused Disease. Int J Mol Sci 19:3426. 10.3390/IJMS19113426

8. Lewis GF, Rader DJ (2005) New insights into the regulation of HDL metabolism and reverse cholesterol transport. Circ Res 96:1221–1232. 10.1161/01.RES.0000170946.56981.5C

9. Faludi AA, Izar MCO, Saraiva JFK, et al (2017) Atualização da Diretriz Brasileira de Dislipidemias e Prevenção da Aterosclerose – 2017. Arq. Bras. Cardiol. 1–76

10. Steinberg D (2009) The LDL modification hypothesis of atherogenesis: An update. J Lipid Res 50:. 10.1194/JLR.R800087-JLR200

11. Ullrich C, Pirchl M, Humpel C (2010) Hypercholesterolemia in rats impairs the cholinergic system and leads to memory deficits. Mol Cell Neurosci 45:408. 10.1016/J.MCN.2010.08.001

12. Moreira ELG, De Oliveira J, Engel DF, et al (2014) Hypercholesterolemia induces short-term spatial memory impairments in mice: Up-regulation of acetylcholinesterase activity as an early and causal event? J Neural Transm. 10.1007/s00702-013-1107-9

13. Paul R, Borah A (2017) Global loss of acetylcholinesterase activity with mitochondrial complexes inhibition and inflammation in brain of hypercholesterolemic mice. Sci Reports 2017 71 7:1–13. 10.1038/s41598-017-17911-z

14. de Oliveira J, Hort MA, Moreira ELG, et al (2011) Positive correlation between elevated plasma cholesterol levels and cognitive impairments in LDL receptor knockout mice: relevance of cortico-cerebral mitochondrial dysfunction and oxidative stress. Neuroscience 197:99–106. 10.1016/J.NEUROSCIENCE.2011.09.009

15. De Oliveira J, Moreira ELG, Mancini G, et al (2013) Diphenyl diselenide prevents cortico-cerebral mitochondrial dysfunction and oxidative stress induced by hypercholesterolemia in LDL receptor knockout mice. Neurochem Res. 10.1007/s11064-013-1110-4

16. Paul R, Choudhury A, Chandra Boruah D, et al (2017) Hypercholesterolemia causes psychomotor abnormalities in mice and alterations in cortico-striatal biogenic amine neurotransmitters: Relevance to Parkinson’s disease. Neurochem Int 108:15–26. 10.1016/J.NEUINT.2017.01.021

17. Prasanthi JRP, Dasari B, Marwarha G, et al (2010) Caffeine protects against oxidative stress and Alzheimer’s disease-like pathology in rabbit hippocampus induced by cholesterol-enriched diet. Free Radic Biol Med 49:1212–1220. 10.1016/J.FREERADBIOMED.2010.07.007

18. Paul R, Choudhury A, Kumar S, et al (2017) Cholesterol contributes to dopamine-neuronal loss in MPTP mouse model of Parkinson’s disease: Involvement of mitochondrial dysfunctions and oxidative stress. PLoS One 12:. 10.1371/JOURNAL.PONE.0171285

19. Björkhem I, Meaney S, Fogelman AM (2004) Brain Cholesterol: Long Secret Life behind a Barrier. Arterioscler. Thromb. Vasc. Biol.

20. Dietschy JM, Turley SD (2004) Cholesterol metabolism in the central nervous system during early development and in the mature animal. J. Lipid Res.

21. Nieweg K, Schaller H, Pfrieger FW (2009) Marked differences in cholesterol synthesis between neurons and glial cells from postnatal rats. J Neurochem 109:125–134. 10.1111/J.1471-4159.2009.05917.X

22. de Oliveira J, Engel DF, de Paula GC, et al (2020) High Cholesterol Diet Exacerbates Blood-Brain Barrier Disruption in LDLr–/– Mice: Impact on Cognitive Function. J Alzheimer’s Dis 78:97–115. 10.3233/JAD-200541

23. Chen X, Ghribi O, Geiger JD (2010) Caffeine protects against disruptions of the blood-brain barrier in animal models of Alzheimer’s and Parkinson’s diseases. In: Journal of Alzheimer’s Disease

24. Thirumangalakudi L, Prakasam A, Zhang R, et al (2008) High cholesterol-induced neuroinflammation and amyloid precursor protein processing correlate with loss of working memory in mice. J Neurochem. 10.1111/j.1471-4159.2008.05415.x

25. Chen YL, Wang LM, Chen Y, et al (2016) Changes in astrocyte functional markers and β-amyloid metabolism-related proteins in the early stages of hypercholesterolemia. Neuroscience 316:178–191. 10.1016/J.NEUROSCIENCE.2015.12.039

26. Rapp JH, Pan XM, Neumann M, et al (2008) Microemboli composed of cholesterol crystals disrupt the blood-brain barrier and reduce cognition. Stroke 39:2354–2361. 10.1161/STROKEAHA.107.496737

27. Chen X, Wagener JF, Morgan DH, et al (2010) Endolysosome mechanisms associated with Alzheimer’s disease-like pathology in rabbits ingesting cholesterol-enriched diet. J Alzheimers Dis 22:1289–1303. 10.3233/JAD-2010-101323

28. Engel DF, de Oliveira J, Lopes JB, et al (2016) Is there an association between hypercholesterolemia and depression? Behavioral evidence from the LDLr-/-mouse experimental model. Behav Brain Res 311:31–38. 10.1016/J.BBR.2016.05.029

29. Hui L, Chen X, Geiger JD (2012) Endolysosome involvement in LDL cholesterol-induced Alzheimer’s disease-like pathology in primary cultured neurons. Life Sci 91:1159–1168. 10.1016/J.LFS.2012.04.039

30. Chen H, Chan DC (2006) Critical dependence of neurons on mitochondrial dynamics. Curr Opin Cell Biol 18:453–459. 10.1016/J.CEB.2006.06.004

31. Knott AB, Bossy-Wetzel E (2008) Impairing the mitochondrial fission and fusion balance: a new mechanism of neurodegeneration. Ann N Y Acad Sci 1147:283–292. 10.1196/ANNALS.1427.030

32. Knott AB, Perkins G, Schwarzenbacher R, Bossy-Wetzel E (2008) Mitochondrial fragmentation in neurodegeneration. Nat Rev Neurosci 9:505–518. 10.1038/NRN2417

33. Rangaraju V, Calloway N, Ryan TA (2014) Activity-driven local ATP synthesis is required for synaptic function. Cell 156:825–835. 10.1016/J.CELL.2013.12.042

34. Freeman LR, Keller JN (2012) Oxidative stress and cerebral endothelial cells: Regulation of the blood-brain-barrier and antioxidant based interventions. Biochim. Biophys. Acta - Mol. Basis Dis.

35. Halliwell B (2011) Free radicals and antioxidants - Quo vadis? Trends Pharmacol. Sci.

36. Pernas L, Scorrano L (2016) Mito-Morphosis: Mitochondrial Fusion, Fission, and Cristae Remodeling as Key Mediators of Cellular Function. Annu Rev Physiol 78:505–531. 10.1146/ANNUREV-PHYSIOL-021115-105011/1

37. Lee H, Yoon Y (2016) Mitochondrial fission and fusion. Biochem Soc Trans 44:1725–1735. 10.1042/BST20160129

38. Frank S, Gaume B, Bergmann-Leitner ES, et al (2001) The role of dynamin-related protein 1, a mediator of mitochondrial fission, in apoptosis. Dev Cell 1:515–525. 10.1016/S1534-5807(01)00055-7

39. Bem AF de, Farina M, Portella R de L, et al (2008) Diphenyl diselenide, a simple glutathione peroxidase mimetic, inhibits human LDL oxidation in vitro. Atherosclerosis 201:92–100. 10.1016/j.atherosclerosis.2008.02.030

40. Lowry OH, Rosebrough NJ, Farr AL, Randall RJ (1951) Protein measurement with the Folin phenol reagent. J Biol Chem. 10.1016/s0021-9258(19)52451-6

41. Farias HR, Gabriel JR, Cecconi ML, et al (2020) The metabolic effect of α-ketoisocaproic acid: in vivo and in vitro studies. Metab Brain Dis. 10.1007/s11011-020-00626-y

42. Cassina A, Radi R (1996) Differential inhibitory action of nitric oxide and peroxynitrite on mitochondrial electron transport. Arch Biochem Biophys. 10.1006/abbi.1996.0178

43. Fischer JC, Ruitenbeek W, Berden JA, et al (1985) Differential investigation of the capacity of succinate oxidation in human skeletal muscle. Clin Chim Acta 153:23–36. 10.1016/0009-8981(85)90135-4

44. Agnello M, Morici G, Rinaldi AM (2008) A method for measuring mitochondrial mass and activity. Cytotechnology 56:145–149. 10.1007/S10616-008-9143-2/FIGURES/3

45. Yan HM, Ramachandran A, Bajt ML, et al (2010) The Oxygen Tension Modulates Acetaminophen-Induced Mitochondrial Oxidant Stress and Cell Injury in Cultured Hepatocytes. Toxicol Sci 117:515. 10.1093/TOXSCI/KFQ208

46. Santos L, Behrens L, Barbosa C, et al (2024) Histone 3 Trimethylation Patterns are Associated with Resilience or Stress Susceptibility in a Rat Model of Major Depression Disorder. Mol Neurobiol 1–20. 10.1007/S12035-024-03912-3/METRICS

47. Halliwell B (2001) Role of free radicals in the neurodegenerative diseases: Therapeutic implications for antioxidant treatment. Drugs and Aging

48. Pernas L, Scorrano L (2016) Mito-Morphosis: Mitochondrial Fusion, Fission, and Cristae Remodeling as Key Mediators of Cellular Function. Annu Rev Physiol 78:505–531. 10.1146/ANNUREV-PHYSIOL-021115-105011/1

49. de Oliveira J, Engel DF, de Paula GC, et al (2020) LDL Receptor Deficiency Does not Alter Brain Amyloid-β Levels but Causes an Exacerbation of Apoptosis. J Alzheimer’s Dis 73:585–596. 10.3233/JAD-190742

50. Diaz G, Batetta B, Sanna F, et al (2008) Lipid droplet changes in proliferating and quiescent 3T3 fibroblasts. Histochem Cell Biol 129:611–621. 10.1007/S00418-008-0402-2/FIGURES/6

51. Greenspan P, Fowler SD (1985) Spectrofluorometric studies of the lipid probe, nile red. J Lipid Res 26:781–789. 10.1016/S0022-2275(20)34307-8

52. Schönfeld P, Reiser G (2013) Why does brain metabolism not favor burning of fatty acids to provide energy-Reflections on disadvantages of the use of free fatty acids as fuel for brain. J Cereb Blood Flow Metab 33:1493–1499. 10.1038/JCBFM.2013.128

53. Yang DS, Stavrides P, Saito M, et al (2014) Defective macroautophagic turnover of brain lipids in the TgCRND8Alzheimer mouse model: prevention by correcting lysosomal proteolyticdeficits. Brain 137:3300. 10.1093/BRAIN/AWU278

54. Kim I, DeBartolo D, Ramanan S, et al (2015) Excess Lipid Accumulation in Cortical Neurons in Multiple Sclerosis May Lead to Autophagic Dysfunction and Neurodegeneration (P5.237). Neurology 84:. 10.1212/WNL.84.14_SUPPLEMENT.P5.237

55. Farmer BC, Walsh AE, Kluemper JC, Johnson LA (2020) Lipid Droplets in Neurodegenerative Disorders. Front Neurosci 14:. 10.3389/FNINS.2020.00742

56. Colebc NB, Murphy DD, Grider T, et al (2002) Lipid droplet binding and oligomerization properties of the Parkinson’s disease protein α-synuclein. J Biol Chem 277:6344–6352. 10.1074/jbc.M108414200

57. Li D, Xu N, Hou Y, et al (2022) Abnormal lipid droplets accumulation induced cognitive deficits in obstructive sleep apnea syndrome mice via JNK/SREBP/ACC pathway but not through PDP1/PDC pathway. Mol Med 28:1–19. 10.1186/S10020-021-00427-8/FIGURES/10

58. Gu H-F, Li H-Z, Xie X-J, et al (2017) Oxidized low-density lipoprotein induced mouse hippocampal HT-cell damage via promoting the shift from autophagy to apoptosis. CNS Neurosci Ther 23. 10.1111/cns.12680

59. Engel DF, Grzyb AN, de Oliveira J, et al (2019) Impaired adult hippocampal neurogenesis in a mouse model of familial hypercholesterolemia: A role for the LDL receptor and cholesterol metabolism in adult neural precursor cells. Mol Metab 30:1–15. 10.1016/j.molmet.2019.09.002

60. Dias IHK, Mistry J, Fell S, et al Oxidized LDL lipids increase β-amyloid production by SH-SY5Y cells through glutathione depletion and lipid raft formation. 10.1016/j.freeradbiomed.2014.07.012

61. Sugawa M, Ikeda S, Kushima Y, et al (1997) Oxidized low density lipoprotein caused CNS neuron cell death. Brain Res 761:165–172. 10.1016/S0006-8993(97)00468-X

62. Demchenko AP (2013) Beyond annexin V: Fluorescence response of cellular membranes to apoptosis. Cytotechnology 65:157–172. 10.1007/s10616-012-9481-y

63. Kiessling V, Wan C, Tamm LK (2009) Domain coupling in asymmetric lipid bilayers. Biochim Biophys Acta - Biomembr 1788:64–71. 10.1016/J.BBAMEM.2008.09.003

64. Yamaji-Hasegawa A, Tsujimoto M (2006) Asymmetric distribution of phospholipids in biomembranes. Biol Pharm Bull 29:1547–1553. 10.1248/BPB.29.1547

65. Bevers EM, Williamson PL (2016) Getting to the outer leaflet: Physiology of phosphatidylserine exposure at the plasma membrane. Physiol Rev 96:605–645. 10.1152/PHYSREV.00020.2015/ASSET/IMAGES/LARGE/Z9J0021627550010.JPEG

66. Elliott MR, Ravichandran KS (2010) Clearance of apoptotic cells: implications in health and disease. J Cell Biol 189:1059–1070. 10.1083/JCB.201004096

67. Segawa K, Nagata S (2015) An Apoptotic “Eat Me” Signal: Phosphatidylserine Exposure. Trends Cell Biol 25:639–650. 10.1016/J.TCB.2015.08.003

68. Kann O, Kovács R, Kann O (2007) Mitochondria and neuronal activity. Am J Physiol Cell Physiol 292:641–657. 10.1152/ajpcell.00222.2006.-Mitochondria

69. Sazanov LA (2015) A giant molecular proton pump: structure and mechanism of respiratory complex I. 10.1038/nrm3997

70. Akhtar F, Bokhari SRA (2022) Apoptosis. StatPearls

71. Nunnari J, Suomalainen A (2012) Mitochondria: In Sickness and in Health. Cell 148:1145. 10.1016/J.CELL.2012.02.035

72. Cottet-Rousselle C, Ronot X, Leverve X, Mayol J-F (2011) Cytometric Assessment of Mitochondria Using Fluorescent Probes. 10.1002/cyto.a.21061

73. Puleston D (2015) Detection of mitochondrial mass, damage, and reactive oxygen species by flow cytometry. Cold Spring Harb Protoc 2015:830–834. 10.1101/PDB.PROT086298

74. Connolly NM, Theurey P, Adam-Vizi V, et al (2017) Guidelines on experimental methods to assess mitochondrial dysfunction in cellular models of neurodegenerative diseases. Cell Death Differ 25:542–572. 10.1038/s41418-017-0020-4

75. Kroemer G, Galluzzi L, Brenner C (2007) Mitochondrial membrane permeabilization in cell death. Physiol Rev 87:99–163. 10.1152/PHYSREV.00013.2006/ASSET/IMAGES/LARGE/Z9J0010724230012.JPEG

76. Sesso A, Belizário JE, Marques MM, et al (2012) Mitochondrial swelling and incipient outer membrane rupture in preapoptotic and apoptotic cells. Anat Rec (Hoboken) 295:1647–1659. 10.1002/AR.22553

77. Wani AA, Rangrez AY, Kumar H, et al (2008) Analysis of reactive oxygen species and antioxidant defenses in complex I deficient patients revealed a specific increase in superoxide dismutase activity. Free Radic Res 42:415–427. 10.1080/10715760802068571

78. Blacker TS, Duchen MR (2016) Investigating mitochondrial redox state using NADH and NADPH autofluorescence. Free Radic Biol Med 100:53–65. 10.1016/J.FREERADBIOMED.2016.08.010

79. Lin MT, Beal MF (2006) Mitochondrial dysfunction and oxidative stress in neurodegenerative diseases. Nature 443:787–795. 10.1038/NATURE05292

80. Wang L, Duan Q, Wang T, et al (2015) Mitochondrial Respiratory Chain Inhibitors Involved in ROS Production Induced by Acute High Concentrations of Iodide and the Effects of SOD as a Protective Factor. 10.1155/2015/217670

81. Keller JN, Hanni KB, Gabbita SP, et al (1999) Oxidized lipoproteins increase reactive oxygen species formation in microglia and astrocyte cell lines. Brain Res 830:10–15. 10.1016/S0006-8993(99)01272-X

82. Montilla P, Espejo I, Muñoz MC, et al (2006) Protective effect of red wine on oxidative stress and antioxidant enzyme activities in the brain and kidney induced by feeding high cholesterol in rats. Clin Nutr 25:146–153. 10.1016/j.clnu.2005.10.004

83. Afonso MS, De O Silva AM, Carvalho EB, et al (2013) Phenolic compounds from Rosemary (Rosmarinus officinalis L.) attenuate oxidative stress and reduce blood cholesterol concentrations in diet-induced hypercholesterolemic rats. Nutr Metab (Lond) 10:19. 10.1186/1743-7075-10-19

84. Otunola GA, Oloyede OB, Oladiji AT, Afolayan AJ (2014) Selected spices and their combination modulate hypercholesterolemia-induced oxidative stress in experimental rats

85. Ridker PM, Everett BM, Thuren T, et al (2017) Antiinflammatory Therapy with Canakinumab for Atherosclerotic Disease. N Engl J Med 377:1119–1131. 10.1056/NEJMoa1707914

86. Cid-Castro C, Hernández-Espinosa DR, Morán J (2018) ROS as Regulators of Mitochondrial Dynamics in Neurons. Cell Mol Neurobiol 2018 385 38:995–1007. 10.1007/S10571-018-0584-7

87. Shutt T, Geoffrion M, Milne R, McBride HM (2012) The intracellular redox state is a core determinant of mitochondrial fusion. EMBO Rep 13:909. 10.1038/EMBOR.2012.128

88. Lacombe A, Scorrano L (2024) The interplay between mitochondrial dynamics and autophagy: From a key homeostatic mechanism to a driver of pathology. Semin Cell Dev Biol 161–162:1–19. 10.1016/J.SEMCDB.2024.02.001

89. Ikeda Y, Shirakabe A, Brady C, et al (2015) Molecular Mechanisms Mediating Mitochondrial Dynamics and Mitophagy and Their Functional Roles in the Cardiovascular System. J Mol Cell Cardiol 0:116. 10.1016/J.YJMCC.2014.09.019

